# Subcellular thermal profiling enables the deep functional exploration of the mitochondrial proteome

**DOI:** 10.1101/2024.02.27.582308

**Authors:** Pablo Rivera-Mejías, Cécile Le Sueur, Nils Kurzawa, Isabelle Becher, Mikhail M Savitski

## Abstract

Mitochondria are membrane-bound organelle hubs of cellular metabolism and signaling. The dysregulation of mitochondria is related to the genesis of several highly prevalent diseases, including cancer and cardiovascular disorders, urging the development of novel technologies to systematically study this organelle and its dynamics. Thermal proteome profiling (TPP) allows the unbiased study of the interactions of proteins with drugs, metabolites, and other proteins, providing a unique understanding of the state of the proteome. Here, we develop and introduce an optimized TPP workflow, mito-TPP, for the direct and extensive study of this organelle. We demonstrate that our approach detects both direct mitochondrial small molecule-protein and metabolite-protein interactions, as well as indirect downstream effects. We also show that mito-TPP preserves features from whole-cell systems, such as the coaggregation of interacting proteins. Finally, we explore the mitochondrial proteoform map, detecting more than 180 proteins with multiple proteoform groups. Overall, we demonstrate that mito-TPP is a powerful new tool for the functional study of the mitochondrial proteome.

## Introduction

Mitochondria are membrane-bound organelles that regulate metabolism and signaling essential to sustaining cellular homeostasis. Dysregulation of mitochondrial proteins is related to the genesis and progression of several highly prevalent disorders, including cancer^1^, neurodegenerative diseases^2^, and cardiovascular disorders^3,4^. Mutations in nuclear-encoded and mitochondrial-encoded genes of mitochondrial proteins are the most common forms of inherited metabolic genetic disorders, giving rise to a spectrum of symptoms, including optic atrophy, muscle weakness, and neurodegeneration^5^. Additionally, mitochondrial proteins are the target of several signaling metabolites in physiological and pathological conditions^6–9^, and they are gaining significant interest as targets for small molecules treating diseases such as cancer^10,11^. This complex regulation of mitochondrial proteins reflects the necessity for developing novel tools for the unbiased study of the mitochondrial proteome.

Thermal proteome profiling (TPP)^12^ measures the system-wide thermal stability of proteins by combining multiplexed mass spectrometry (MS)-based quantitative proteomics^13,14^ with the principle of the cellular thermal shift assay^15,16^. Thermal stability reports the tolerance of a protein to heat-induced denaturation and aggregation, a phenomenon that depends on the context and molecular interactions of the protein^15^. The quantification of the soluble fraction of a sample across a heat gradient allows the generation of melting curves of all detectable proteins^17^. TPP enables the detection of the targets of small molecules^12,18,19^, protein-metabolite interactions^20–22^, and protein-protein interactions^15,23^. It has also been used to study the effect of post-translational modifications (PTM)^24–26^ and different cellular states, like cell cycle progression^27,28^ on protein thermal stability. Finally, this method has been recently applied for the detection of functional proteoform groups^29^, where proteins encoded by the same gene but differing in their final form (e.g. due to alternative splicing, proteolytic cleavage, or PTMs^30^) can have distinct functions. This versatility reflects the power of TPP for exploring the state of the proteome beyond changes in protein abundance, and the potential of this method for studying mitochondria.

Current protocols are designed to apply TPP to whole cells^31,32^. Although this has been very useful, the direct application of TPP over isolated mitochondria would bring several experimental advantages, enabling the study of mitochondria-specific and direct effects of perturbations, and the definition of organelle-specific functions of mitochondrial proteins that can also be localized in other cellular compartments^33^. Furthermore, this workflow would make it possible to use molecules with low plasma membrane permeability, as is the case for some metabolites, since the compound could be delivered directly to the organelle. Finally, analysis of the isolated mitochondrial sub-proteome would increase the approach’s sensitivity, as key but low-abundant mitochondrial proteins can be challenging to identify by MS when performing the standard TPP workflow on whole cells.

Here, we develop a modified TPP approach, mito-TPP, which ultimately enables the direct application of TPP on isolated mitochondria. Overall, we demonstrate that mito-TPP facilitates the systematic exploration of direct interactions of mitochondrial proteins with small molecules and metabolites, protein-protein interactions, and the exploration of the mitochondrial proteoform map.

## Results

### Development of mito-TPP

To examine the effect of performing a TPP on isolated mitochondria, we applied a temperature range over crude mitochondrial fractions obtained by differential centrifugation (Fig. 1). After removing aggregates by filtration, we labeled the soluble protein fractions with tandem mass tags (TMT) and subjected them to MS analysis, enabling the multiplexed relative quantification of proteins across different temperatures^17^. We focused our analysis on high-confidence mitochondrial proteins based on the MitoCarta3.0 database^34^. We observed that mitochondrial isolation impacts the melting profiles of mitochondrial proteins (Supplementary Fig. 1a) compared to whole-cell lysates (Supplementary Fig. 1b). Specifically, this is manifested in a lack of aggregation of proteins at higher temperatures (Supplementary Fig. 1c), which would prohibit detecting thermal stability changes at these temperatures, limiting the usefulness of the method.

**Fig. 1:**
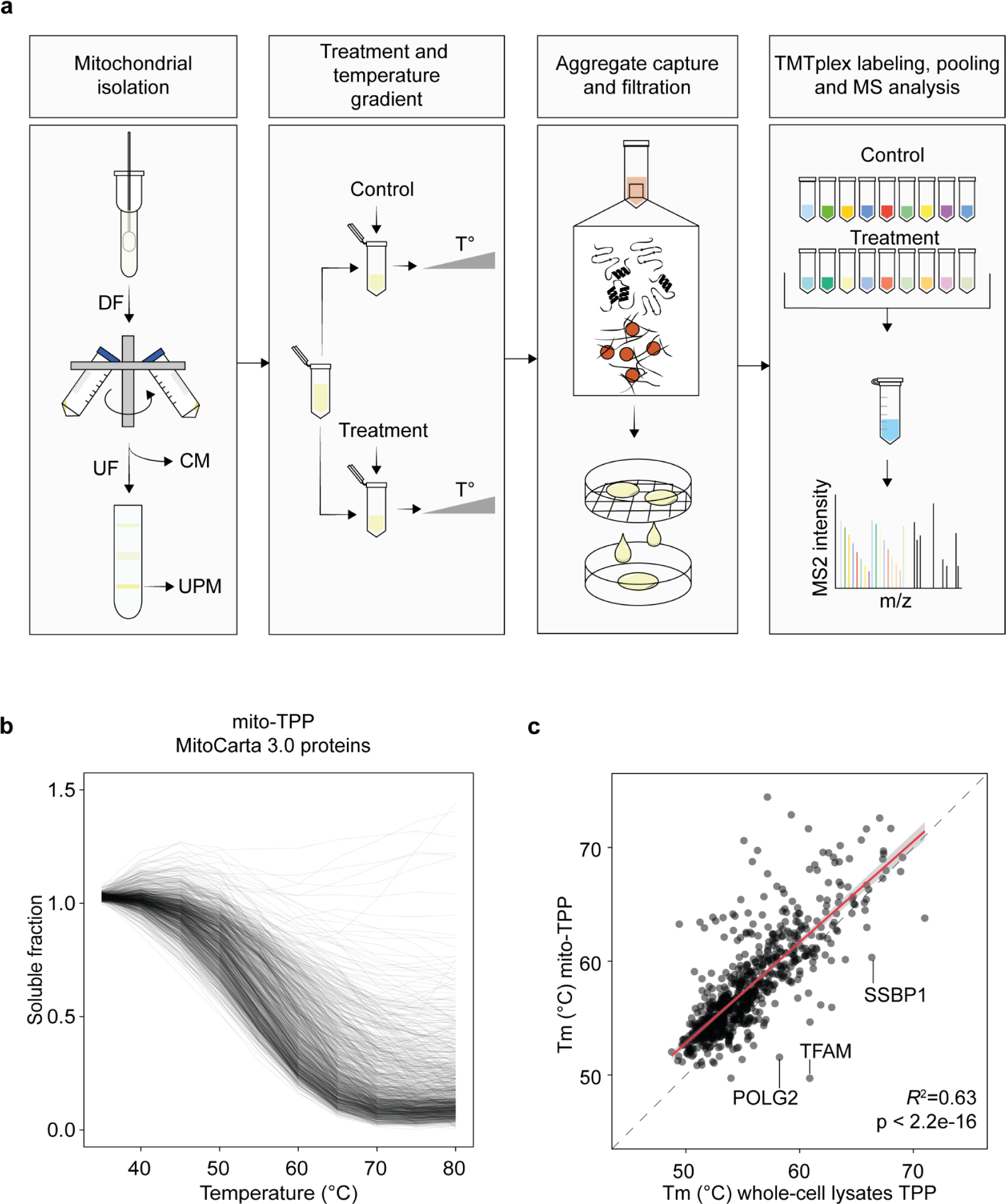
Subcellular thermal profiling of mitochondria. **a**, Mitochondrial isolation is performed by differential centrifugation (DF) to obtain crude mitochondrial fractions (CM), followed by density gradient ultracentrifugation (UC) to obtain ultra-pure mitochondria (UPM). In the presence or absence of a compound, isolated mitochondria are treated with a heat gradient, followed by aggregate capture with microparticles, lysis, and aggregate filtration. Soluble proteins are digested with trypsin, and peptide mixtures are analyzed by multiplexed mass spectrometry-based quantitative proteomics. **b**, Melting curves of mitochondrial proteins (MitoCarta3.0) obtained with mito-TPP. Data are fitted splines (df = 5, n = 1). **c**, Scatterplot comparing protein Tm values determined from whole-cell lysates or mito-TPP. The red linear regression curve shows the agreement between the datasets (*R*^2^ =0.63, p-value >2.2e-16).

We hypothesized that the lack of aggregation is due to the removal, during the purification step, of proteins that would normally promote aggregation by acting as aggregation seeds and/or enhancing molecular crowding. Magnetic microparticles used for standard MS-based proteomics sample preparation can capture and stabilize protein aggregates induced by different perturbations, including heating^35^. Therefore, we tested if adding microparticles to our workflow would improve the protein melting profiles in isolated mitochondria. After heating, we applied hydrophobic magnetic beads at different beads-to-proteins ratios (1:4, 1:2, 1:1, and 2:1 µg/µg), observing a clear improvement in the protein melting profiles, including at the lower ratios (Supplementary Fig. 1d and Supplementary Fig. 1e). Thus, our workflow incorporates both mitochondrial enrichment and protein aggregates stabilization steps (Fig. 1a).

To evaluate changes in the thermal stability of mitochondrial proteins, we compared the melting points (Tm, the temperature at which the protein soluble fraction is reduced by 50%) obtained with this workflow to those obtained in whole-cell lysates (Fig. 1b and Supplementary Fig. 1b). On the one hand, we observed an overall increase (2.3 °C on average) in Tm after performing a TPP in isolated mitochondria compared to whole-cell lysates (Supplementary Fig. 2a), affecting all mitochondrial sub-compartments to a similar extent, except for the intermembrane space (IMS) (Supplementary Fig. 2b). On the other hand, only a few proteins showed decreased Tm, including TFAM, POLG2, and SSBP1, proteins involved in mitochondrial DNA (mtDNA) homeostasis^36^ (Fig. 1c), indicating a loss of interactions required for the stability of these proteins during the purification steps. Aside from the systematic shift in melting behavior, we obtained a significant agreement between Tms of isolated mitochondria and whole-cell lysates (Fig. 1c, *R*^2^=0.63, p < 2.2e-16), together with the aggregation of the vast majority of mitochondrial proteins in the purified mitochondria at higher temperatures (Supplementary Fig. 1c). Importantly, we quantified a higher number of peptides (10787 versus 8234), proteins (899 versus 812, unique peptides > 1), and significantly higher sequence coverage (Supplementary Fig. 2d) in the isolated mitochondria compared to whole-cell lysates. These results were reached by measuring 12 hours for isolated mitochondria compared to 24 hours for whole-cell lysates, decreasing the instrument time by half per experiment while significantly gaining in mitochondrial proteome coverage. Taken together, we adapted the application of TPP to isolated mitochondria, termed mito-TPP (Fig. 1a), by compensating for the loss of cellular crowding and reaching a deeper coverage of the mitochondrial proteome.

### Studying the effects of Antimycin A on the mitochondrial proteome with mito-TPP

To assess if mito-TPP can detect small molecule-protein interactions, we first treated isolated mitochondria with Antimycin A (AA), which binds the complex III of the electron transport chain (ETC)^37^. We incubated the samples with AA for 15 minutes at room temperature, followed by heating with a higher temperature range (40-70°C) than the one commonly used in mammalian cells in order to accommodate the increase in thermal stability observed in the mito-TPP setup. Following this, we applied our recently developed hierarchical Gaussian processes model (GP-Melt) method to analyze the protein melting curves^38^. GP-Melt allows the study of non-sigmoidal and non-melter proteins, together with the scaling of the normalized intensities with the mean value of the protein per replicate, a scaling value more robust to technical errors than using the lowest tested temperature^38^. Finally, we also calculated the area between the curves (ABC) to measure the effect size of thermal stability changes, where positive ABC values reflect stabilization and negative values show the destabilization of the protein in the presence of the treatment.

We focused our analysis on proteins with significant stability changes (adjusted p-value < 0.05) and an effect size cut off of |ABC| > 3. Using these criteria, we observed significant changes in 305 mitochondrial proteins (out of 893 detected in at least two replicates), out of which AA destabilized 299 and stabilized 6 proteins. Among the 6 stabilized mitochondrial proteins, 5 are complex III subunits, including the respiratory subunit CYC1, the core subunits UQCRC1 and UQCRC2, and the small subunit UQCRQ ^39^(Fig. 2a, b). Across the other detected subunits, we also observed the stabilization of the small subunit UQCR10 (Supplementary Fig. 3a). The mitochondrially encoded protein MT-CYB, which directly binds AA at the Qi site^40^, did not pass our filter for protein identification (unique peptides > 1). However, we observed reproducible stabilization of this unique peptide across all three replicates (Supplementary Fig. 3b). We did not observe changes in the thermal stability of the other detected complex III subunits UQCRFS1, UQCRH, and UQCRB (Supplementary Fig. 3a). UQCRH and UQCRB are small subunits localized in the IMS and mitochondrial matrix^39^. Importantly, these two proteins do not co-melt with the rest of the complex III subunits (Supplementary Fig. 5c), helping to explain their different response to AA.

**Fig. 2:**
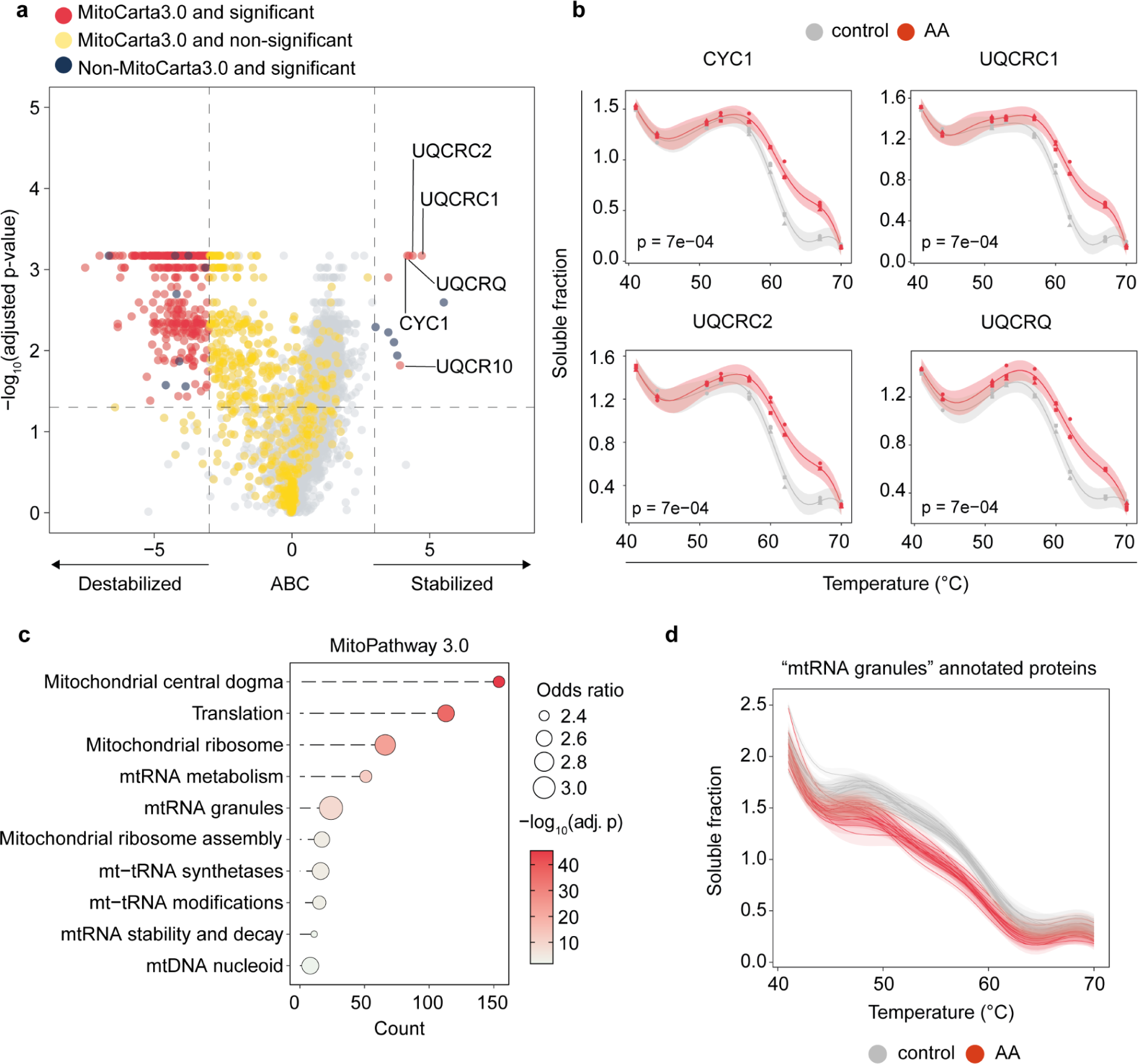
Assessing the effect of Antimycin A in the mitochondrial proteome with mito-TPP. **a**, Volcano plot comparing the effect size (ABC = area between the curves) and statistical significance (adjusted p-value) of crude mitochondrial fractions treated with 5 µM AA. Dots are colored depending on their annotation in the MitoCarta3.0 list and their significance (significative changed protein = adjusted p-value < 0.05 and |ABC| > 3). Vertical dashed lines indicate ABC thresholds and the horizontal line indicates the adjusted p-value threshold. **b**, Melting curves of complex III subunits in the presence or absence of AA 5 µM. Data are the mean melting curve plus 95% confidence regions (CR). p = adjusted p-value. **c**, Dot plot of overrepresented MitoPathway 3.0 terms (q value < 0.05) among mitochondrial proteins destabilized by AA treatment. **d**, Melting curves of significantly changed proteins by AA annotated as “mt-RNA granules” members. Data are the mean melting curve plus 95% CR per protein. p = adjusted p-value (n = 3).

The inhibition of complex III by AA decreases the mitochondrial membrane potential (MMP)^41^, an electrochemical property that drives several biochemical reactions within this organelle^42^. Importantly, the loss of MMP also disrupts the mitochondrial matrix structure, leading to a swollen morphology^43–45^. We observed that the majority of the destabilized proteins are localized in the mitochondrial matrix (Supplementary Fig. 3c). In this compartment, more than 40% of the annotated proteins are affected by AA. Therefore, these results suggest that the broad AA-induced destabilization of mitochondrial proteins is linked to the loss of MMP. Finally, we observed an enrichment of pathways associated with the mitochondrial ribosome and mitochondrial RNA (mtRNA) metabolism across the destabilized proteins (Fig. 2c), including components of the mtRNA granules (Fig. 2d), a pathway not previously related to MPP loss.

Altogether, our data demonstrated that mito-TPP enables the study of direct interactions of small molecules with mitochondrial proteins and complexes and the downstream effects caused by these interactions.

### Study of protein-metabolite interactions with mito-TPP

To test the ability of mito-TPP to identify protein-metabolite interactions, we incubated isolated mitochondria with pyruvate, a crucial metabolite for mitochondrial metabolism. Pyruvate is imported into the matrix by the mitochondrial pyruvate carrier (MPC) and metabolized to fuel in the tricarboxylic acid (TCA) cycle^46–48^. To prevent confounding effects that may arise from downstream metabolites generated from pyruvate, we incubated isolated mitochondria only for 10 minutes at room temperature. Additionally, we took advantage of the loss of matrix metabolites and cofactors due to the long sequential centrifugation steps and the basal activity of metabolite carriers^49,50^, which should prevent the conversion of pyruvate into its products.

Upon the treatment with pyruvate, we detected significant changes in thermal stability for just 19 mitochondrial proteins (out of 889 detected in at least two replicates) (adjusted p-value < 0.05, |ABC| > 3), contrary to the broad effects observed with AA. Remarkably, we observed the thermal stabilization of the two subunits of the MPC (MPC1 and MPC2) among the top stabilized hits (Fig. 3a, b), highlighting the sensitivity of our method to detect the interaction between carriers and their metabolites. This effect was highly specific as no other detected mitochondrial metabolite carriers (55 annotated as small molecule transporters) were stabilized by pyruvate. We also observed the stabilization of the matrix proteins BCAT2 and OAT among the top stabilized hits (Fig. 3c, d). These aminotransferases are part of the branched-chain amino acid^51^ and ornithine metabolism pathways^52^, respectively, whose interaction with pyruvate has not been previously reported. Other mitochondrial enzymes with aminotransferase activity, ABAT and GOT2, and the cytosolic enzymes detected in our crude mitochondrial fractions, BCAT1 and GOT1, were also significantly stabilized (adjusted p-value < 0.05) (Supplementary Fig. 4), suggesting a possible general crosstalk between aminotransferase enzymes and pyruvate. We also detected the stabilization of HMGCL (Fig. 3a), an enzyme that participates in ketogenesis^53^ and the last step of leucine catabolism^54^, supporting once again the crosstalk of pyruvate and branched-chain amino acid catabolism. Previous studies also support this interaction^55–57^. We did not detect significant changes in TCA cycle-related enzymes, in line with the idea that our experimental design prevents downstream metabolic reactions, strengthening the conclusion that the observed effects are due to the direct action of pyruvate.

**Fig. 3:**
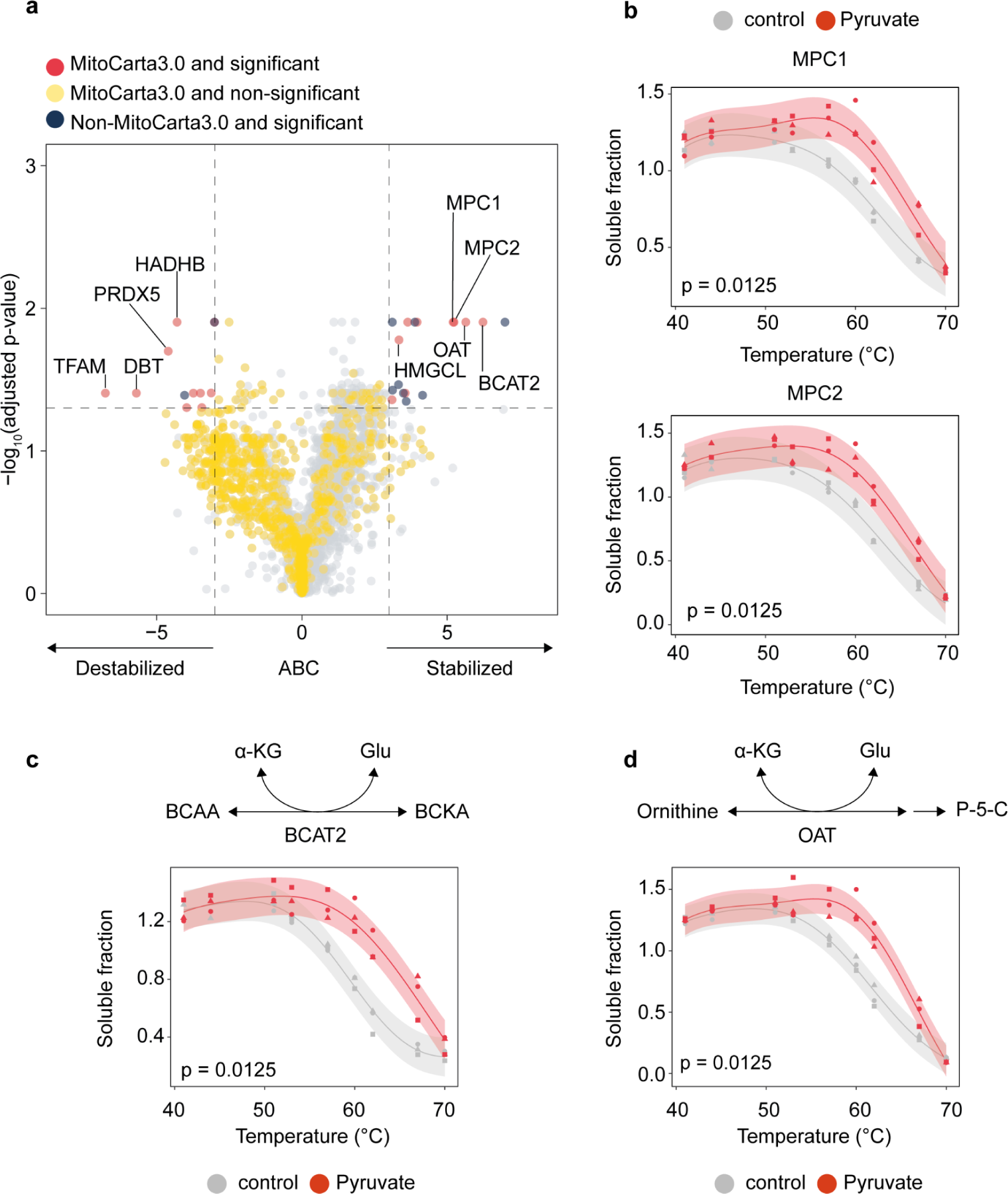
Searching protein pyruvate interactors with mito-TPP. **a**, Volcano plot comparing the effect size (ABC = area between the curves) and statistical significance (adjusted p-value) of crude mitochondrial fractions treated with 5 mM Pyruvate. Dots are colored depending on their annotation in the MitoCarta3.0 list and their significance (significative changed protein = adjusted p-value < 0.05 and |ABC| > 3). Vertical dashed lines indicate ABC thresholds and the horizontal line indicates the adjusted p-value threshold. **b**, Melting curves of mitochondrial pyruvate carrier (MPC) subunits in the presence or absence of pyruvate 5 mM. Melting curve of BCAT2 (**c**) and OAT (**d**) in the presence or absence of pyruvate 5 mM. Data are the mean melting curve plus 95% CR. p = adjusted p-value (n = 3).

In conclusion, mito-TPP can detect protein-metabolite interactions in mitochondria. Remarkably, our data also showed the potential of mito-TPP for studying mitochondrial metabolite carriers.

### Profiling of protein-protein interactions with mito-TPP

TPP can also be used to study protein complexes in cells based on the observation that interacting proteins show similar melting profiles^23^. Thermal proximity coaggregation analysis (TPCA) calculates the similarity or distance between melting curves of proteins to assess the presence or absence of an interaction. This method has been used to explore protein complex dynamics in response to drug treatment^23^, cell cycle stages^28^, viral infection^58,59^, and T-cell activation^60^.

Thus, to test if we can assess the coaggregation of protein complexes with mito-TPP, we applied our workflow to ultra-pure mitochondrial samples obtained by differential centrifugation followed by a Percoll density gradient and ultracentrifugation (Fig. 1a). Using this approach, we obtained a higher enrichment of the mitochondrial proteome compared to whole-cell lysates and crude mitochondrial fractions (Supplementary Fig. 2c), quantifying 923 mitochondrial proteins present in the three independent replicates.

To evaluate the performance of our method, we first manually annotated known mitochondrial protein complexes using different sources, including the CORUM database^61^ and the MitoPathway annotation^34^. This list includes the electron transport chain, mitochondrial ribosome, protein import, and ultrastructure organization complexes (Supplementary Table 1). Next, we computed the median Euclidean distance between each complex member and compared the probability of observing a similar value across 10.000 random complexes. Finally, we assessed the ability of our workflow to predict co-melting of true complexes by calculating a receiver operating characteristic (ROC) curve, observing an area under the curve (AUC) of 0.74 (Supplementary Fig. 5a). The magnitude of the AUC value is similar to the one observed for whole-cell complexes from intact cells^23^, supporting that mito-TPP preserves the coaggregation properties of protein complexes, which enables a more comprehensive exploration of the mitochondrial proteome.

Across the complexes with significant nonrandom co-melting of their subunits, we observed the large and small mitochondrial ribosomes (Fig. 4a). Most of the proteins belonging to this complex showed a similar melting profile except MRPS36 (gray melting curve). This protein was previously described as a component of the small ribosomal subunit^62,63^. However, MRPS36 was not located in the structure of the human mitochondrial ribosome^64^. Additionally, other studies demonstrated that MRPS36 is a member of the oxoglutarate dehydrogenase complex, mediating the interaction between the E2o (DLST) and E3 (DLD) subunits, as shown by complexome profiling and XL-MS^65–68^. In line with this, we observed a similarity between the melting curves of MRPS36, DLST, and DLD (Supplementary Fig. 5b). Therefore, our data provide additional evidence to support that MRPS36 is not a subunit of the mitochondrial ribosome and highlight the complementarity of our method with other techniques for exploring mitochondrial protein complexes.

**Fig. 4:**
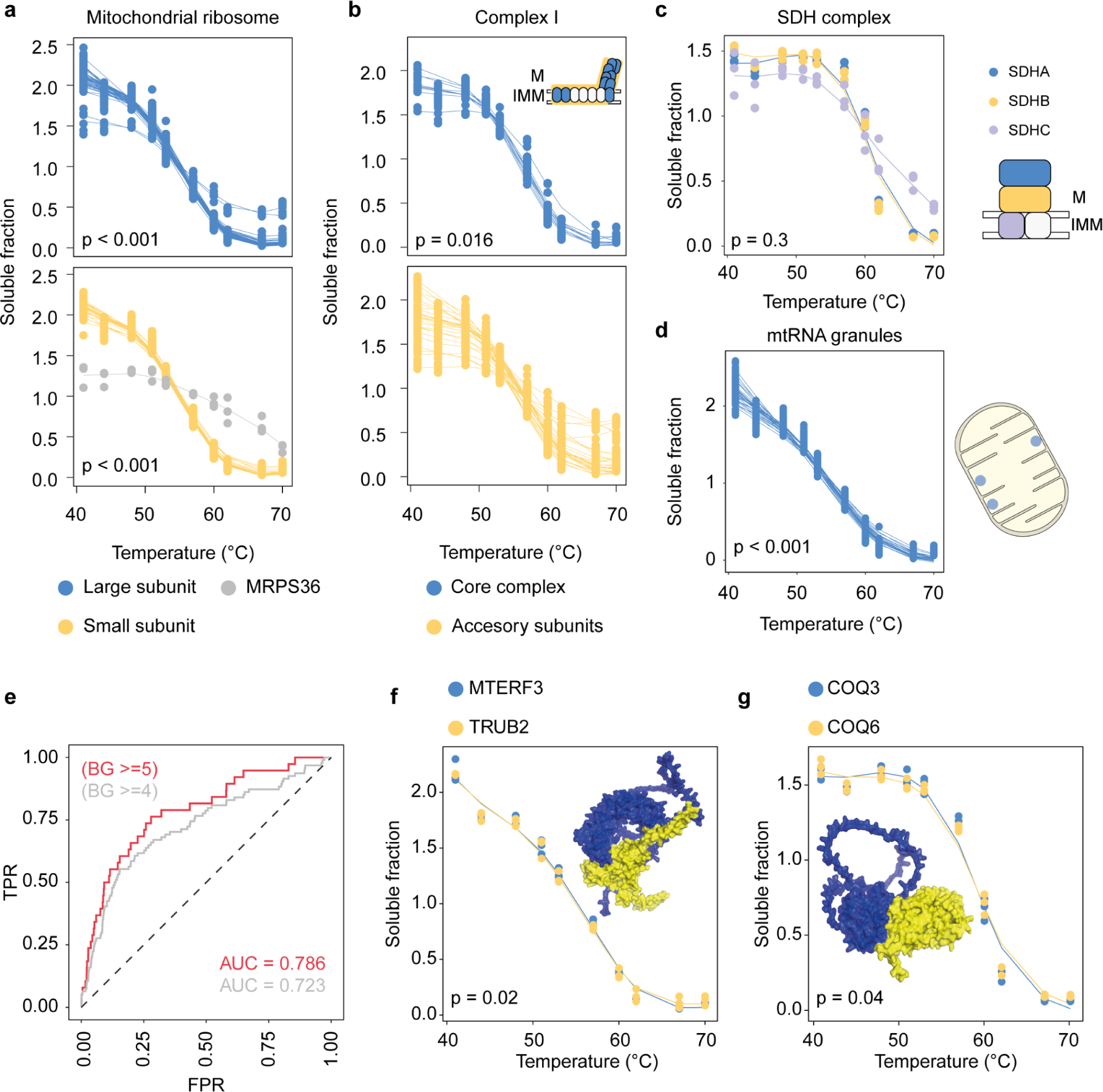
Mitochondrial thermal proximity coaggregation profiling. Melting curves of large and small mitochondrial ribosome subunits (**a**), Core and accessory complex I subunits (adjusted p-value for whole complex coaggregation) (**b**), complex II subunits (**c**), and mtRNA granules proteins (**d**). For complex I (**b**), the model represents the spatial distribution of core (blue) and accessory subunits (yellow). For complex II (**c**), the model represents the spatial distribution of the matrix subunits (blue and yellow), compared to the membrane subunits (purple). White colored subunits represent not detected proteins. IMM = inner mitochondrial membrane; M = mitochondrial matrix. The model in (**d**), represents the sub-mitochondrial distribution of mtRNA granules, attached to the IMM^45^. Data are fitted splines (df = 5). p = adjusted p-value (n = 3). **e**, Receiver operating characteristic (ROC) curve of recovered PPI of mitochondrial proteins reported in the BioGRID database. FPR = false positive rate; TPR = true positive rate; AUC = area under the curve; BG = BioGRID count. **f**, **g**, melting curves of the indicated PPI pairs. AlphaFold2 models were obtained from Pei.et al. 2002^73^. Data are fitted splines (df = 5). p = adjusted p-value (n = 3).

Interestingly, along with finding a significant nonrandom coaggregation of all complex I subunits (adjusted p-value = 0.016), we observed that core complex proteins melt more similarly than the accessory subunits, suggesting that the coaggregation pattern of mitochondrial proteins can resemble their spatial distribution within the complex (Fig. 4b). Similarly, the two complex II subunits localized in the matrix, SDHA and SDHB, showed a similar melting compared to the membrane protein SDHC (Fig. 4c), leading to a non-significant co-melting of the whole complex (p = 0.3). In addition, UQCRH and UQCRB showed a different melting behavior compared to the rest of complex III (Supplementary Fig. 5c). As described above, UQCRH and UQCRB are two small subunits exposed to the IMS and the mitochondrial matrix^39^, respectively, also unaffected by AA (Supplementary Fig. 3a). Taken together, we conclude that the coaggregation of mitochondrial proteins is influenced by their spatial distribution within the complex, as previously shown for bacteria^69^.

Finally, we found that proteins belonging to mtRNA granules also showed a significant nonrandom coaggregation (Fig. 4d). This observation is supported by their co-destabilization in the presence of AA (Fig. 2d), revealing that coaggregation is also a property of proteins located in membrane-less condensates.

We also tested the performance of our method for predicting protein-protein interaction (PPI) pairs by using BioGRID^70^ annotations of mitochondrial proteins, discarding homodimers (Supplementary Table 2). Like with protein complexes, our approach showed a good performance in detecting true PPI pairs, particularly when increasing the evidence supporting the interaction (BG, BioGRID count) (Fig. 4e).

Next, we used a recent data set that explores PPI from mitochondria using AlphaFold2^71^ and RoseTTAFold^72^ models to explore unknown PPI or PPI with low experimental evidence^73^. We focused our analysis on protein pairs with high contact probability for AlphaFold2 and RoseTTAFold models (>= 0.9). Interestingly, across the significant co-melting pairs, we identified the interaction between TRUB2 and MTERF3, a PPI with low experimental evidence (BG < 4). On the one hand, TRUB2 is a pseudouridine synthase annotated as a component of mtRNA granules^74^. On the other hand, MTERF3 regulates mitochondrial ribosome biogenesis and is necessary for normal mitochondrial transcription^75,76^. This interaction was previously reported by proximity biotinylation and co-migration in a 2D gel electrophoresis^74^, showing a contact probability score higher than 0.99 in the *in silico* models. Our data also supports this direct interaction, observing a significant nonrandom co-melting (adjusted p-value= 0.02) for this protein pair (Fig. 4f). Furthermore, our data also supports the direct interaction between COQ3 and COQ6 (adjusted p-value = 0.04) (Fig. 4g), two components of the coenzymes Q biosynthetic pathway^77^, not previously reported to interact directly but with a high contact probability score according to the AlphaFold2 and RoseTTAFold models (>0.99). These data demonstrate that our workflow can be combined with additional experimental and *in silico* approaches for exploring protein-protein interactions within mitochondria.

### Exploration of mitochondrial proteoforms with mito-TPP

The current maps of the mitochondrial proteome comprise approximately 1130 proteins (MitoCarta3.0^34^, MitoCop^78^), annotated as a unique entity per gene. However, it is well documented that from a single gene, a diversity of proteoforms can arise due to alternative splicing events, single amino-acid polymorphism, post-translational modifications, and proteolytic cleavage^30,79^. Notably, proteoforms generated from the same gene can differ in their activities^80^, highlighting the importance of the identification and functional characterization of these proteoforms.

Recently, an approach combining TPP with deep fractionation for high sequence coverage was developed to detect functional proteoform groups ^29^. We applied this method to the ultra-pure mitochondrial data by computing pairwise similarities between melting curves of peptides mapped to the same gene, followed by the construction of similarity networks and community detection for the assignment of proteoform groups (Fig. 5a). As we described above, the higher purity of these fractions allows to obtain a higher enrichment of mitochondrial proteins (Supplementary Fig. 2c) and a higher protein coverage (Fig. 5b and Supplementary Fig. 2d) compared to whole-cell and crude mitochondrial isolates, quantifying 13201 peptides from mitochondrial proteins across the three replicates. These characteristics enabled the exploration of proteoforms specifically localized in mitochondria.

**Figure 5.**
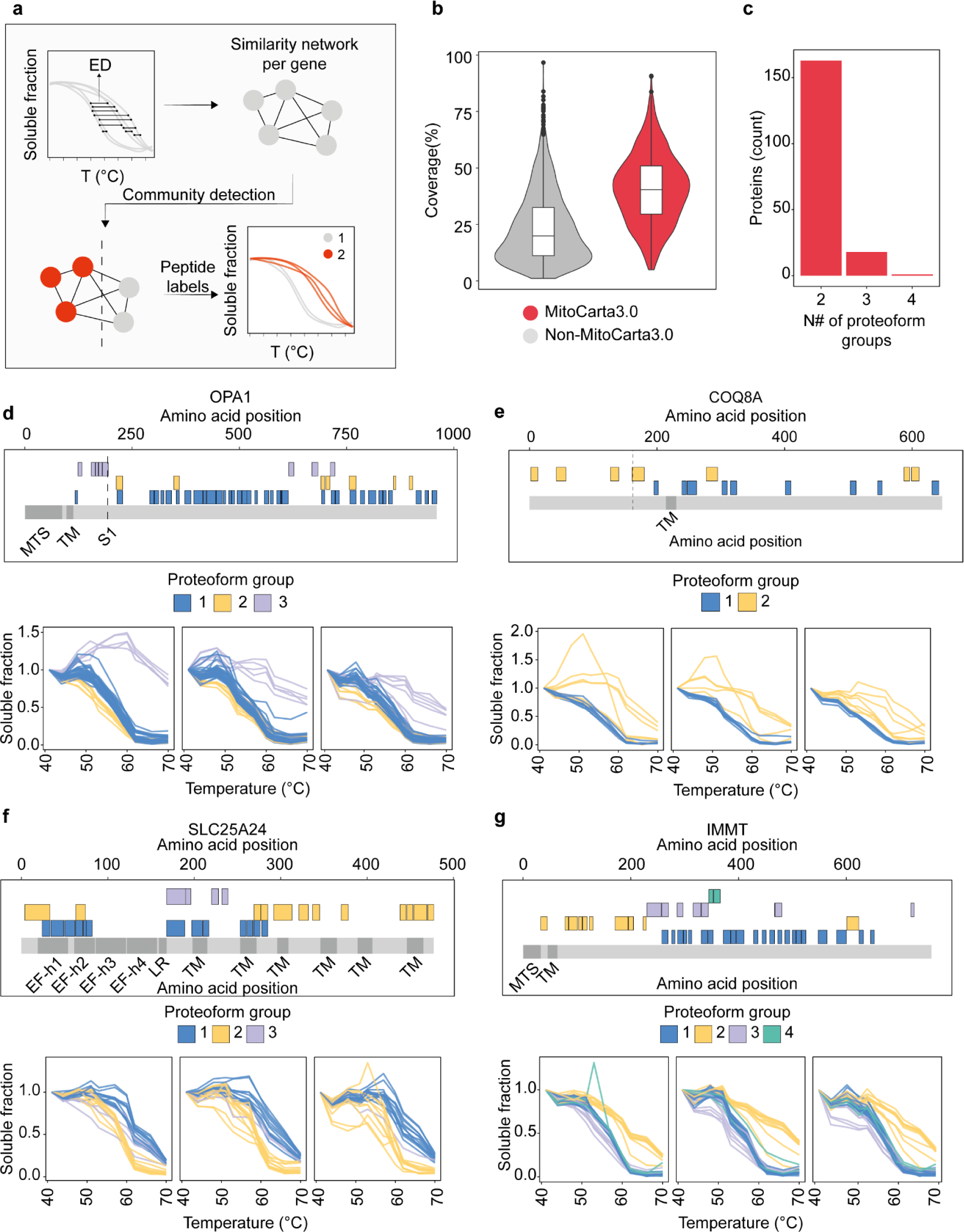
Detection of functional proteoform groups using up-mito-TPP. **a**, Workflow for the detection of functional proteoform groups. **b**, Average coverage (%) of mitochondrial proteins across three replicates, compared to non-mitochondrial proteins detected in the samples. **c**, Bar plot of the number of identified proteoform groups of mitochondrial proteins with multiple proteoforms. **c**-**g**, peptide localization across the protein amino acid sequence (upper panels, x-axis = amino acid position) and melting curves (lower panels) of those peptides of the indicated proteins, colored by their proteoform group. Data are connected dots per peptide. MTS = mitochondrial targeting sequence; TM = transmembrane domain; EF-h = EF-hand domain; LR = linker region. (n=3).

From the 923 mitochondrial proteins (MitoCarta3.0) detected in the three independent replicates, we identified 182 proteins with more than one proteoform group, out of which 163 had 2, 13 had 3, and one had 4 groups (Fig. 5c and Supplementary Table 3). We did not observe any particular preferential sub-mitochondrial localization of proteins with multiple proteoforms (Supplementary Fig. 6a).

Among the detected proteoforms, we found exciting examples of previously known, others that strengthen previous literature claims, and completely new mitochondrial proteoforms. OPA1 is a membrane-anchored protein localized in the IMM, whose mutation causes autosomal dominant optic atrophy^81^. This protein presents several transmembrane isoforms (long forms, L-OPA1) generated by alternative splicing^82^, which can be subjected to proteolytic cleavage to generate soluble short forms (S-OPA1)^83^. Long and short forms are present in steady-state conditions and differ in their functions^84,85^. We observed a proteoform group with higher thermal stability (Fig. 5d), where most of its peptides localized before the S1 cleavage site of OPA1 (group 3). Therefore, we assigned the membrane-anchored L-OPA1 forms to the proteoform 3 and the other two proteoform groups to the S-OPA1 forms.

Similarly, we detected two proteoform groups for COQ8A (Fig. 5e). COQ8A is an enzyme essential for synthesizing coenzyme Q, whose mutation causes cerebellar ataxia^86,87^. Recent reports showed that COQ8A is processed to generate the mature form of this protein, lacking the last 162 amino acids^88,89^. We observed a proteoform group with higher thermal stability, with several peptides located before the proposed cleavage site. Hence, our data supports the presence of short and long COQ8A forms, which correlate to the mature and immature states, respectively. However, we cannot discard that the long form could also play a functional role in mitochondria since the requirement of the proteolytic cleavage for the correct function of COQ8A has not been tested.

SLC25A24 is a solute carrier that transports Mg-ATP to the matrix in exchange for phosphate. Mutations in this carrier are associated with the development of human progeroid Fontaine syndrome^90^. SLC25A24 is composed of two main regions. The transmembrane region mediates the transport of substrates, and the N-terminal regulatory region with two pairs of EF-hands domains regulates the activity of the carrier in response to calcium^91^. In the absence of calcium, the regulatory region interacts with the transmembrane, preventing substrate exchange in a close conformation. This interaction is released to an open state in the presence of calcium, allowing substrate transport^92,93^.

Interestingly, a study showed that the calcium-bound, open conformation of SLC25A24, has lower thermal stability than the closed state^93^. Here, we identified two main proteoform groups with different thermal stability (Fig. 5f). This result indicates that detected proteoforms could reflect the different conformational states of the carrier. Importantly, our assay buffer does not contain chelators, giving the possibility of retaining a calcium pool bound to proteins. Although further experiments are required to test this model, this data points out the potential of our workflow for the study of the functional impact of disease mutation or modification in mitochondrial solute carriers.

Finally, we identified IMMT as the protein with most proteoform groups (4) in our data set. IMMT is a MICOS complex protein essential for maintaining crista junctions and the contacts between the inner and outer mitochondrial membrane^94^. Splice forms of this protein are described in the Uniprot database and recent studies in human induced pluripotent stem cell-derived cardiomyocytes^95^. However, no evidence exists of IMMT proteoforms at the protein level. We identified four different proteoform groups for IMMT, showing different thermal stability (Fig. 5g). Immunoblotting experiments supported this finding, detecting several bands around the expected molecular weight (Supplementary Fig. 6b), showing different melting behavior, particularly the bands located at lower molecular weights, resembling the observations at the peptide level.

Together, we demonstrated the utility of mito-TPP for the exploration of known and new mitochondrial proteoforms, achieving the expansion of the current mitochondrial proteome maps.

## Discussion

Here, we present a workflow for the subcellular thermal profiling of mitochondria. The mito-TPP approach allows a faster and deeper study of the direct effects of small molecules and metabolites in mitochondrial proteins, including metabolite carriers. This method also enables a deeper exploration of mitochondrial PPIs and the detection of mitochondrial proteoform groups.

As described before, the isolation of mitochondria by differential centrifugation leads to the loss of matrix metabolites^49,50^. On the one hand, this can be exploited to avoid further conversion of the target metabolite into the downstream products, preventing possible confounding effects. On the other hand, this could also lead to a loss of interactions that depend on additional cofactors or metabolites. For example, we did not detect significant changes in the thermal stability of pyruvate dehydrogenase kinases, which are described to interact with pyruvate. However, this interaction requires the presence of ADP^96–99^, which we did not use in the present work. Therefore, future experiments could be performed in the presence of specific metabolite pools, depending on the tested hypothesis. In summary, our method will enable testing the effect of drugs and metabolites directly on mitochondrial proteins in intact isolated mitochondria, including compounds with low plasma-membrane permeability or solubility, such as nucleotides, TCA cycle metabolites, or lipids.

We also demonstrate the utility of mito-TPP for exploring mitochondrial protein complexes. Previous studies showed the loss of the coaggregation patterns of PPIs in cell lysates compared to intact cells. Nevertheless, we observed that those patterns remain in our approach, which we explain by the membrane-bound nature of mitochondria, and highlights the integrity of our samples. Our results also revealed the complementarity of our method with other *in vitro* and *in silico* techniques for the exploration of mitochondrial PPIs.

Here, we also take the first steps towards understanding the mitochondrial proteoform diversity, identifying around 180 mitochondrial proteins with more than one functional proteoform group. It is important to note that the detection of proteoforms using TPP helps to identify and detect functional differences between them since variations in their thermal stability reflect their biophysical properties. For example, the membrane-anchored L-OPA1 isoforms generated by alternative splicing do not differ in function. However, the functional proteoform difference arises from its proteolytic cleavage, generating the soluble S-OPA1 form. We observed that this distinction is reflected in the melting profiles, emphasizing the utility of our workflow for the study of mitochondrial proteoforms. Therefore, our data is a rich resource for further experimental characterization of the functional proteoform diversity.

We anticipate that mito-TPP will help understand the molecular mechanisms underlying pathologies associated with mitochondrial dysregulation and develop therapies to treat those conditions. Additionally, we expect that our workflow principles will facilitate the application of TPP to other subcellular fractions and organelles.

## Methods

### Reagents and cell culture

HEK293T cells were cultured in a DMEM medium (Sigma-Aldrich, D5648), supplemented with 4.6 g/L glucose and 10% fetal calf serum (Gibco, 10270). Cells were cultured at 37°C and 5% CO_2_ and were routinely tested for Mycoplasma infections.

### Crude and ultra-pure mitochondrial purification

Cells were plated 4 days before the experiments until reaching ∼90% confluency. All the subsequent steps were performed at 4°C. Cells were collected by scraping, washed twice with cold PBS, pelleted at 600 g for 5 minutes, and resuspended with mitochondrial isolation buffer (MIB) containing 220 mM mannitol, 70 mM sucrose, and 20 mM HEPES-KOH pH 7.4. Cells were homogenized with 20 strokes using a glass Teflon homogenizer at 1000 rpm. Cell homogenates were centrifuged at 600 g for 5 minutes. The supernatant was collected in a second tube, and the pellet resuspended in MIB and subjected to 10 strokes at 1000 rpm. This second cell homogenate was centrifuged at 600 g for 5 minutes, and the supernatant was mixed with the first one and centrifuged at 1000 g for 5 minutes. The resulting supernatant was centrifuged twice at 10000 g for 10 minutes, and the pellet was resuspended in MIB buffer, corresponding to the crude mitochondrial fraction. For ultra-pure mitochondrial purification, crude mitochondria were subsequently purified on a Percoll (GE Healthcare) density gradient of 12%-19%-40%, centrifuged at 42000 g for 30 minutes using an SW40 rotor in a Beckman Coulter Optima L-100 XP ultracentrifuge. Ultra-pure mitochondria were collected from the third layer, located between 19% and 40%, and washed three times with MIB buffer, centrifuging at 16000 g for 5 minutes.

### Thermal proteome profiling of isolated mitochondria

For studying the effect of metabolites or drugs on mitochondrial proteins, crude mitochondria suspensions were treated with 5 µM Antimycin A (Sigma Aldrich, cat#A8674) or 5 mM sodium pyruvate (Gibco, cat#11360), shaking at 300 RPM at 25°C. Then, nine 20 µL aliquots of the mitochondrial suspensions (3.5 µg/µL final protein concentration) were transferred into a 96-well PCR plate, centrifuged at 100 g for 30 seconds, and heated for 5 minutes (40°-70°C) in a PCR machine (Agilent SureCycler 8800). Next, the samples were placed at room temperature, and 20 uL of Sera-Mag^TM^ SpeedBead Carboxilate-Modified Magnetic Particles suspension (∼0.87 µg/µL) (Cytiva #65152105050250) were added on top of the samples, shacked at 1000 rpm per 10 seconds, and incubated at room temperature for 10 minutes, shaking at 400 rpm. Afterward, the samples were flash-frozen in liquid nitrogen and stored at −70°C until further procedures.

For the coaggregation analysis and functional proteoform detection, 20 µL aliquots of ultra-pure mitochondrial fractions (3.5 µg/µL) were transferred into a 96-well PCR plate and subjected to the same procedure described for crude mitochondrial suspension.

### Thermal proteome profiling of cell lysates

HEK293T cells were collected, washed twice with cold PBS, pelleted at 600 g for 5 minutes, and resuspended in MIB buffer. Then, cells were homogenized with 20 strokes using a glass Teflon homogenizer at 1000 rpm. Then, nine 20 µL aliquots of the cell lysates (3.5 µg/µL final protein concentration) were transferred into a 96-well PCR plate, centrifuged at 100 g for 30 seconds, and heated for 3 minutes (35°-80°C) in a PCR machine (Agilent SureCycler 8800). Next, the samples were placed at room temperature for 3 minutes, flash-frozen in liquid nitrogen, and stored at −70°C until further procedures.

### Lysis

Frozen mitochondrial samples were thawed at 25°C for 1 minute and lysed by adding 20 µL of lysis buffer (0.5% NP-40 final, cOmpleteTM protease inhibitor, and PhosSTOP), resuspending 20 times with a multichannel pipette. Then, the samples were incubated for 30 minutes at 4°C, shaking at 300 rpm. Next, the sample plate was centrifuged for 5 minutes at 4000 g and 4°C. The aggregates were removed from the supernatant using a 0.45 µm pore size 96 well filter plate (Millipore, cat#MSHVN4550). The flow-through was transferred into a new plate, mixed 1:1 with 2x sample buffer (180 mM Tris pH 6.8, 4% SDS, 20% glycerol, 0.1 g bromophenol blue), and stored at −20°C.

For cell lysates, frozen samples were thawed at 25°C for 1 minute and lysed by adding 20 µL of lysis buffer (0.8% NP-40 final, cOmpleteTM protease inhibitor, PhosSTOP, 2.5mM MgCl2, and 0.417U/µL benzonase), resuspending 20 times with a multichannel pipette. Then, the samples were incubated for 1 hour at 4°C, shaking at 400 rpm. Finally, the same procedure for mitochondrial samples was followed.

### Digestion and labeling

The protein samples were digested following a modified SP3 protocol ^100^. In brief, equal volumes of samples across temperatures (equivalent to 5ug of proteins of the lowest temperature) were mixed with water up to reach 20 µL and added to a bead suspension composed by 1:1 mix Sera-Mag A (Cytiva #45152105050250) and Sera-Mag-B (Cytiva #65152105050250), 30 µL ethanol and 10 µL 15% formic acid. This mix was incubated at room temperature for 15 minutes, shaking at 500 rpm. Next, the mix was placed into a 0.45 µm 96 well filter plate, and the bead-bound proteins were washed three times with 70% ethanol. After, 40 µL of digestion solution (1.25 mM TCEP, 5 mM chloroacetamide, 0.2 µg trypsin/Lys-C mix (Promega, cat#V5073), in 100 mM HEPES pH 8.5) were added on top for protein digestion and incubated overnight at room temperature.

The next day, peptides were eluted by centrifugation and vacuum-dried. Next, peptides were resuspended in 10 µL of water and labeled for 1 h at room temperature with 4 µL (∼11 µg/µL in acetonitrile) of TMT10plex or TMT18plex (Thermo Fisher Scientific). The reaction was quenched with 4 µL of 5% hydroxylamine. Experiments belonging to the same biological replicate were combined. Pooled samples were desalted with solid-phase extraction using an OASIS HLB µElution 30 µm plate (Waters, cat#186001828BA), washed twice with 100 µL 0.05% formic acid, eluted with 100 µL 80% acetonitrile, and vacuum dried.

### High-pH fractionation

Dried peptide samples were resuspended in 17 µL 20 mM ammonium formate, pH 10, and fractionated onto 48 fractions using C18-based reversed-phase chromatography with a Phenomenex Gemini 3-µm C18 110-Å 100-mm × 1-mm column. The samples were resolved over an 85 minutes gradient (mobile phase A: 20 mM ammonium formate (pH 10) and mobile phase B: acetonitrile) at 0.1 mL/min starting at 0%B, followed by a linear increase to 35% B from 2 to 60 minutes, with a subsequent increase to 85%B from 62 minutes up to 68 minutes, followed by a linear decrease to 0%B up to 70 minutes, finishing with the equilibration of the system at 0%B until the end of the run. Finally, fractions were concatenated into 6 pools for crude mitochondria experiments and 12 for ultra-pure mitochondria experiments.

### LC-MS

Fractionated peptides were resuspended in 0.05% formic acid and separated using an UltiMate 3000 RSLC nano-LC system (Thermo Fisher Scientific) equipped with an analytical column (Waters nanoEase HSS C18 T3, 75 μm × 25 cm, 1.8 μm, 100 Å) and a trapping cartridge (precolumn; C18 PepMap 100, 5 μm, 300-μm i.d. × 5 mm, 100 Å). The LC system was directly coupled to a Q Exactive Plus mass spectrometer (Thermo Fisher Scientific) using a Nanospray-Flex ion source and a PicoTip Emitter (360-μm o.d. × 20-μm i.d.; 10-μm tip, New Objective). The mobile phase constituted 0.1% formic acid in LC–MS-grade water (buffer A) and 0.1% formic acid in LC–MS-grade acetonitrile (buffer B). Peptides were loaded onto the trapping cartridge using a flow of 30 μL/min of solvent A for 3 minutes and eluted with a constant flow of 0.3 µL/min using 120 minutes of analysis time. During the elution step, the percentage of solvent B was increased stepwise: 2% to 4% B in 4 min, from 4% to 8% in 2 min, 8% to 28% in 96 min, and from 28% to 40% in another 10 min. A column cleaning step using 80% B for 3 min was applied before the system was set again to its initial conditions (2% B) for re-equilibration for 10 minutes. The mass spectrometer was operated in positive ion mode with a spray voltage of 2.2 kV and a capillary temperature of 275 °C. Full-scan mass spectrometry spectra with a mass range of 375–1,200 m/z were acquired in profile mode using a resolution of 70,000 (maximum fill time of 250 ms or a maximum of 3×106 ions (automatic gain control, AGC)). The instrument cycled between MS and MS/MS acquisition in a data-dependent mode and consecutively fragmented the Top 10 peaks with charges 2–4 on the MS scan. MS/MS spectra were acquired in profile mode in the Orbitrap with a resolution of 35,000, a maximum fill time of 120 ms, and an AGC target of 2×105 ions, with a 30-s dynamic exclusion window (normalized collision energy was 30).

### Protein identification and quantification

Mass spectrometry raw files were converted to mzmL format using MSConvert (Proteowizard) using peak picking from the vendor algorithm. Files were searched using MSFragger^101^ v3.8 in Fragpipe v20.0 against a database containing Homo sapiens UniProt FASTA files (proteome ID UP000005640, downloaded on 30 June 2023) along with known contaminants and the reverse protein sequences. The search parameters were as followed: tryptic digestion; maximum missed cleavages = 2; peptide length = 7-50; precursor mass tolerance = 20 ppm, fragment mass tolerance = 20; minimum peaks = 15; topN peaks = 300; fixed modifications were carbamidomethyl on cysteines and TMT10plex or TMT18plex on lysine; variable modifications included acetylation on protein N-terminus, oxidation of methionine, and TMT10plex or TMT18plex on peptide N-termini. Peptide Spectrum Matches (PSMs) validation was performed by Philosopher version 5.1.0, and the false discovery rate was fixed at 1% at the PSMs, peptides, and proteins level. For TMT quantification unique and razor peptides were used. Protein or peptide tsv files were processed using the R programming language.

### TMs calculation

The calculation of Tms for whole-cell lysates and isolated mitochondria was performed as previously described ^17^, using the *TPP* package. In brief, the protein raw intensities were scaled using the lowest tested temperature, and curve fitting and parameter calculations were performed over those values without normalization.

### Data preprocessing and melting analysis for AA and pyruvate experiments

For experiments performed with AA or Pyruvate, we first filtered for proteins detected in at least two replicates and with more than 1 unique peptide. We also discarded the 53°C data from the first replicate of the control condition in the AA experiment due to a technical error. We performed the data preprocessing by aligning the raw intensities across MS runs by selecting the lowest temperature per condition in each experiment and calculating the geometric mean across those values per protein. Second, we computed a scaling value consisting of the raw intensity of the lowest temperature per condition divided by the obtained geometric mean. Third, we aligned all the intensities by multiplying the raw intensities against the scaling value per MS run. Fourth, we normalized the aligned intensities using quantile normalization. Finally, we divided the normalized intensities by the mean of the replicate for scaling.

We utilized the GP-Melt method to analyze melting curves and hit detection, as previously described^38^. In brief, this method models the data through an interpretable hierarchy. The top level of the hierarchy corresponds to the protein combining all conditions and replicates; the second level describes the conditions and the third level accounts for the replicates of each condition. The model fitting gives the user a predictive distribution for the melting curves. For data visualization, the melting curves are represented by their predictive distribution’s mean and 95% Confidence Interval (95% CI). A null model is defined for hits calling, in which the two conditions are merged into a hypothetical unique condition. A Likelihood Ratio test (LRT) value is computed by comparing the likelihood of the alternative and null models. A p-value is derived from the LRT by approximating the null distribution of the statistic. This approximation consists of drawing ten samples per protein according to a null model, fitting the previously defined three-level HGP model on these samples to obtain values of the LRT under the null hypothesis, and combining these values to form the null distribution approximation. Consequently, p-values are defined analogously to permutation p-values. Finally, we adjusted the obtained p-values using the Benhamini-Hochberg (BH) procedure. The ABC was computed from the posterior mean of the predicted distributions.

### Data processing and coaggregation analysis

Data preprocessing for coaggregation analysis was performed as for AA and pyruvate experiments. Then, for the co-melting analysis, we utilized the *Rtpca* package^102^. Briefly, we computed the average Euclidean distance between melting curves of proteins of the same complex (with at least two detected subunits) or PPI pairs, as described in Tan et al (2018)^23^. For the p-value calculation, we estimated the probability of observing the mean Euclidean distance in a distribution of average Euclidean distance values from 10.000 randomly generated complexes or PPI pairs. We adjusted the obtained p-values using the BH procedure. We used an FDR <10% to consider a significant nonrandom comelting. ROC curves and AUC calculations were performed using the same R package. Splines were fitted to the data using the *ggformula* package only for data visualization.

### Data processing and functional proteoform group detection

Detection of proteoform groups from peptide melting profiles was performed as described previously^29^. First, we normalized the peptide intensities across the replicates per temperature, using the variance stabilizing normalization^103^. Then, for proteins with at least ten quantified peptides, similarities of melting profiles of peptides mapping to the same proteins were computed with f(x) = 1 / (1 + d), where d is the Euclidean distance inversely weighted by the number of valid comparisons between two peptide profiles across all samples. Next, a graph was built for each protein with peptides as nodes and similarities as edges and then clustered using the Leiden algorithm^104^ using a resolution of 1.05. The modularity was computed for each protein graph, and only those proteoform groups with a value higher than 10^-13^ were considered further; others were considered as a single proteoform group per protein. Lastly, proteoform groups were merged if their median profile had a Euclidean distance of equal or less than 0.4 to each other. The resolution parameter and the distance threshold were tuned on the protein OPA1 based on prior information on existing proteoforms.

### MitoPathway3.0 over-representation analysis

The over-representation analysis using MitoPathway3.0 annotation^34^, was performed using the *clusterProfilter* package^105^, using all the annotated MitoCarta3.0 proteins as background. p-values were adjusted using the BH procedure, and pathways with a q value <0.05 were considered to be significantly overrepresented.

### Western blotting

Crude mitochondrial samples were heated from 40-70°C, and the soluble fraction was collected as described for mito-TPP. Equal sample volumes were mixed with 4X Laemmli Sample Buffer (Bio-Rad, cat#1910747) plus β-Mercaptoethanol (50 mM). Samples were size-resolved using SDS-PAGE, using a 7.5% Criterion™ TGX™ Precast Midi Protein Gel (Bio-Rad, cat#5671084) and 1× Laemmli buffer. The proteins were transferred onto a 0.2-µm PVDF membrane using semi-dry transfer in the Trans-Blot Turbo Transfer System (Bio-Rad) and 1× Trans-Blot transfer buffer (Bio-rad, cat#10026938). The membrane was blocked with 5% non-fat milk prepared in PBS containing 0.1% Tween. Rabbit polyclonal IgG against IMMT (Proteintech, cat# 10179-1-AP) was diluted (1:1000 dilution) and incubated with the membrane overnight at 4 °C. The membrane was washed three times and incubated with mouse anti-rabbit IgG–HRP (Santa Cruz Biotechnology, cat#sc-2357) for 1 h at room temperature. The membranes were washed and developed using an enhanced chemiluminescence kit (Bio-Rad) using the manufacturer’s instructions.

## Acknowledgments

We thank the members of the Savitski group for the insightful discussions and feedback on the manuscript. We also thank the Proteomics core facility at the EMBL for the assistance with the proteomics experiments. This work was supported by the European Molecular Biology Laboratory. P.R.M. was supported by a Bridging Excellence Fellowship from the EMBL|Stanford Life Science Alliance. M.M.S. is supported by the Allen Distinguished Investigator award through the Paul G. Allen Frontiers Group. C.L.S was supported by a fellowship of the EMBL International PhD programme.

## Authors contributions

P.R.M and M.M.S. conceived and designed the study. P.R.M. performed the experiments together with I.B.. C.L.S. performed the preprocessing and analysis of AA and pyruvate experiments. N.K. performed the preprocessing and analysis for functional proteoform detection, and provided inputs for data interpretation. P.R.M. performed the preprocessing and analysis for TPCA. P.R.M. and M.M.S. drafted the manuscript with contributions from C.L.S., N.K., I.B., and other members from the Savitski lab, which was reviewed and edited by all authors.

## Data availability

The mass spectrometry proteomics data will be deposited to the ProteomeXchange Consortium via the PRIDE partner repository.

---

**Supplementary Figure 1:**
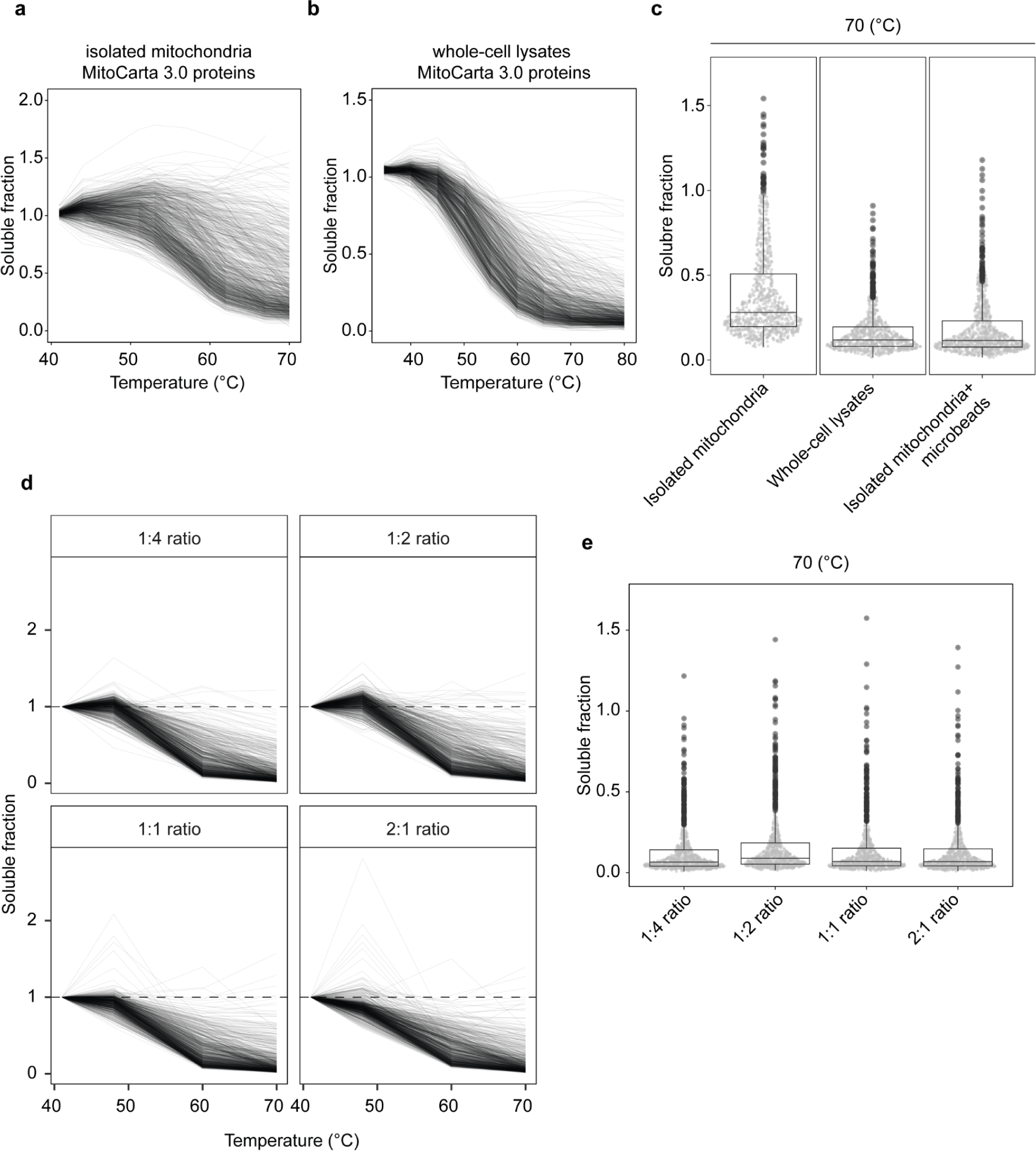
Effect of microparticles on melting profiles of mitochondrial proteins. **a**, Melting curves of mitochondrial proteins (MitoCarta3.0) obtained from isolated mitochondria (n = 1). Data are fitted splines (df = 5). **b**, Melting curves of mitochondrial proteins (MitoCarta3.0) obtained from whole-cell lysates (n = 1). Data are fitted splines (df = 5). **c**, Soluble fraction values of mitochondrial proteins at 70°C obtained from the indicated samples. **d**, Melting curves of mitochondrial proteins (MitoCarta3.0) from isolated mitochondria incubated with the indicated beads-to-protein ratios (µg/ µg) after heating (n = 1). **e**, Soluble fraction of mitochondrial proteins at 70°C incubated with the indicated beads-to-protein ratios.

**Supplementary Figure 2.**
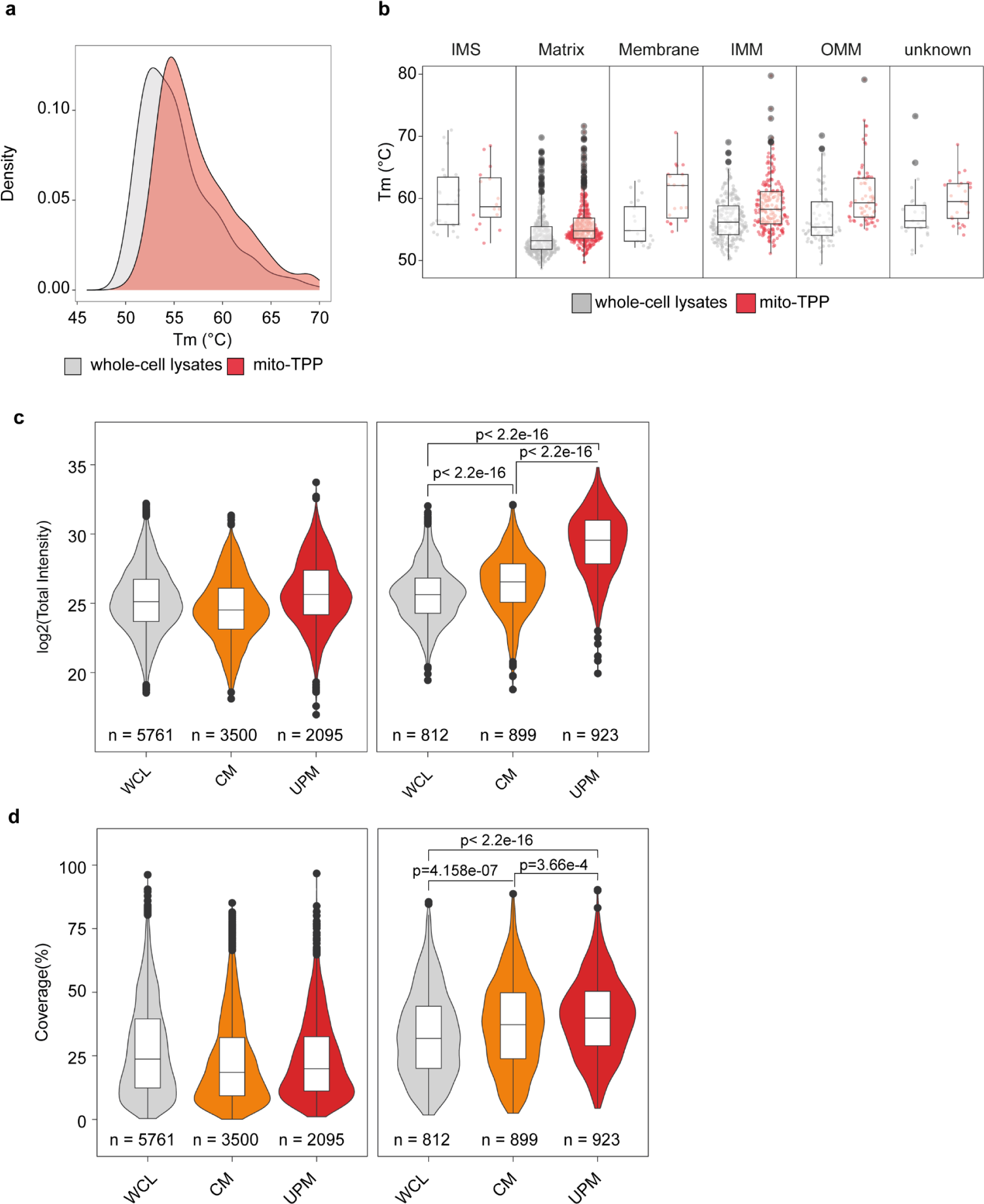
Comparisons between whole-cell and mito-TPP. **a**, Density plot of Tm values of proteins detected in both whole-cell lysates and mito-TPP. **b**, Box plots of Tm values of mitochondrial proteins per sub-mitochondrial localization in whole-cell lysates or mito-TPP. **c**, Enrichment (total intensity) of mitochondrial and non-mitochondrial proteins obtained from the indicated samples. WCL = whole-cell lysates; CM = crude mitochondria (from mito-TPP); UP = ultra-pure mitochondria (the values are the average total intensity per protein across the three independent replicates). n = number of quantified proteins, unique peptides > 1. **d**, Coverage of mitochondrial and non-mitochondrial proteins obtained from the indicated samples. WCL = whole-cell lysates; CM = crude mitochondria (from mito-TPP); UP = ultra-pure mitochondria (the values are the average total intensity per protein across the three independent replicates). Significance (p-values) were calculated with a two-sided Wilcoxon signed-rank test. n = number of quantified proteins, unique peptides > 1.

**Supplementary Figure 3:**
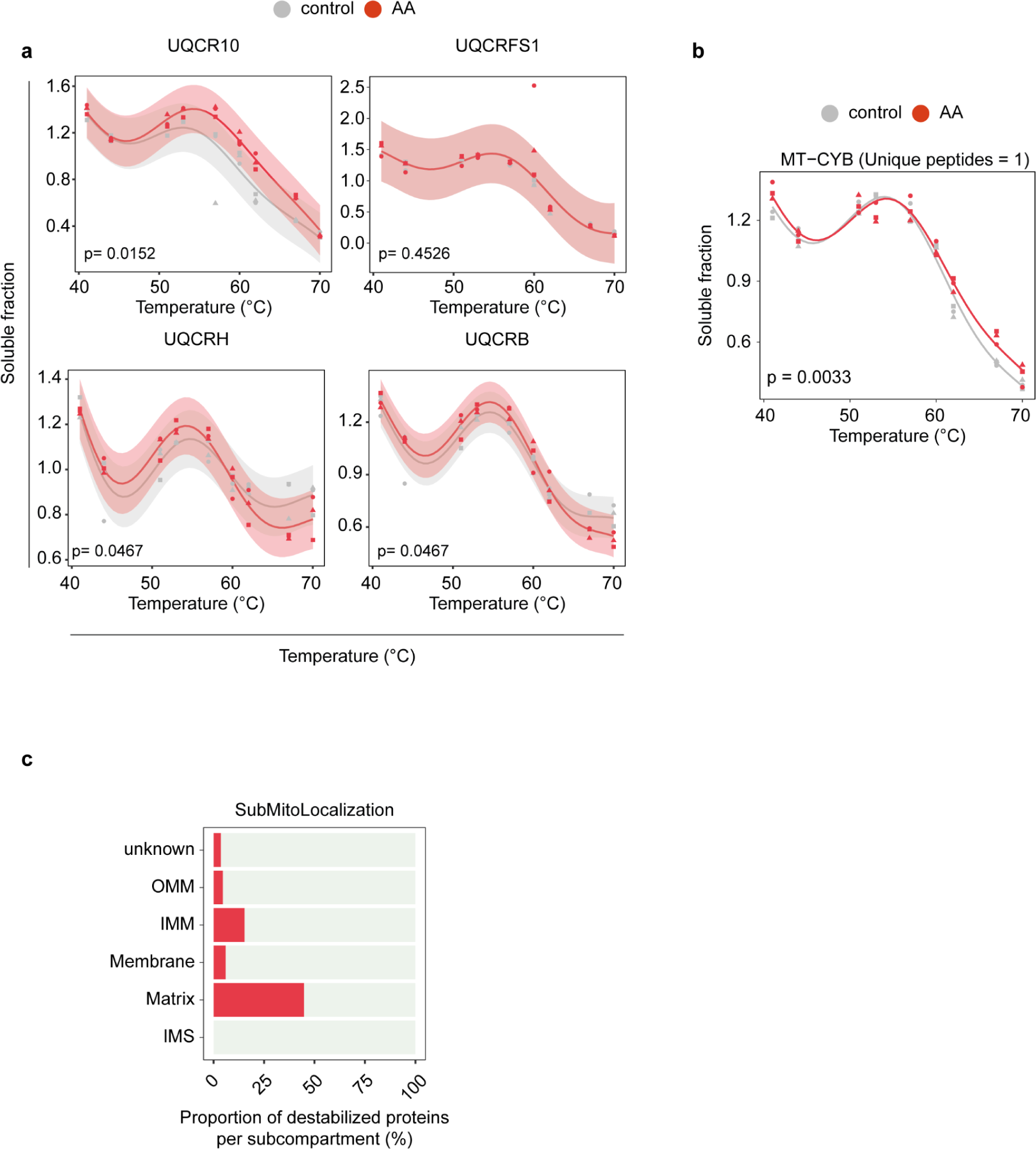
Effects of Antimycin A in the mitochondrial proteome. **a**, Melting curves of complex III subunits in the presence or absence of AA 5 µM. Data are the mean melting curve plus 95% CR per protein. p = adjusted p-value (n = 3).**c**, Melting curves of the peptide detected for MT-CYB (n=3). The indicated p-value was calculated independently. **d**, Sub-mitochondrial localization of destabilized mitochondrial proteins in proportion to all proteins annotated in that subcompartment.

**Supplementary Figure 4:**
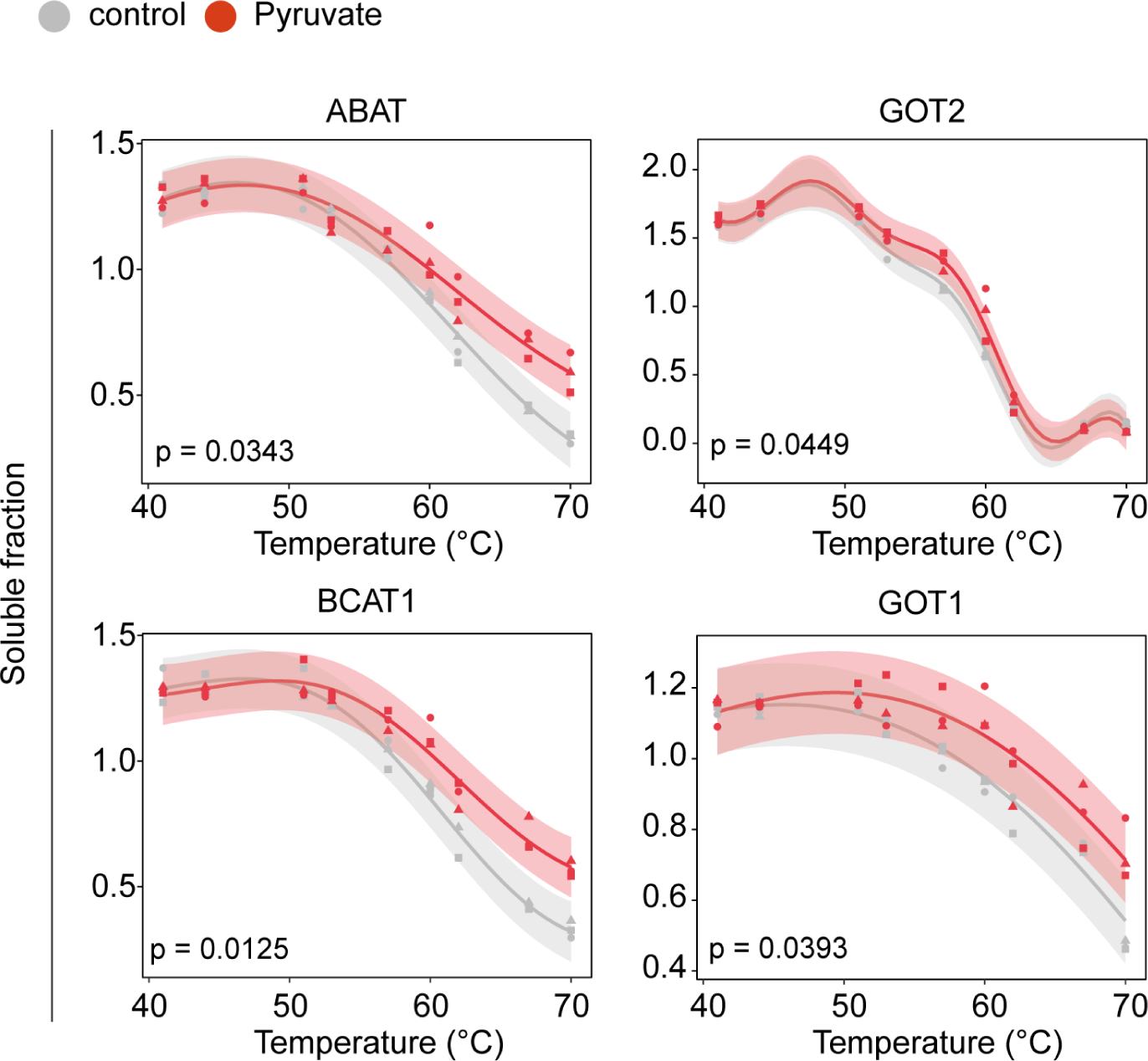
Effect of pyruvate on aminotransferases. Melting curves of the indicated aminotransferases in the presence or absence of 5 mM pyruvate. Data are the mean melting curve plus 95% CR. p = adjusted p-value (n = 3).

**Supplementary Figure 5:**
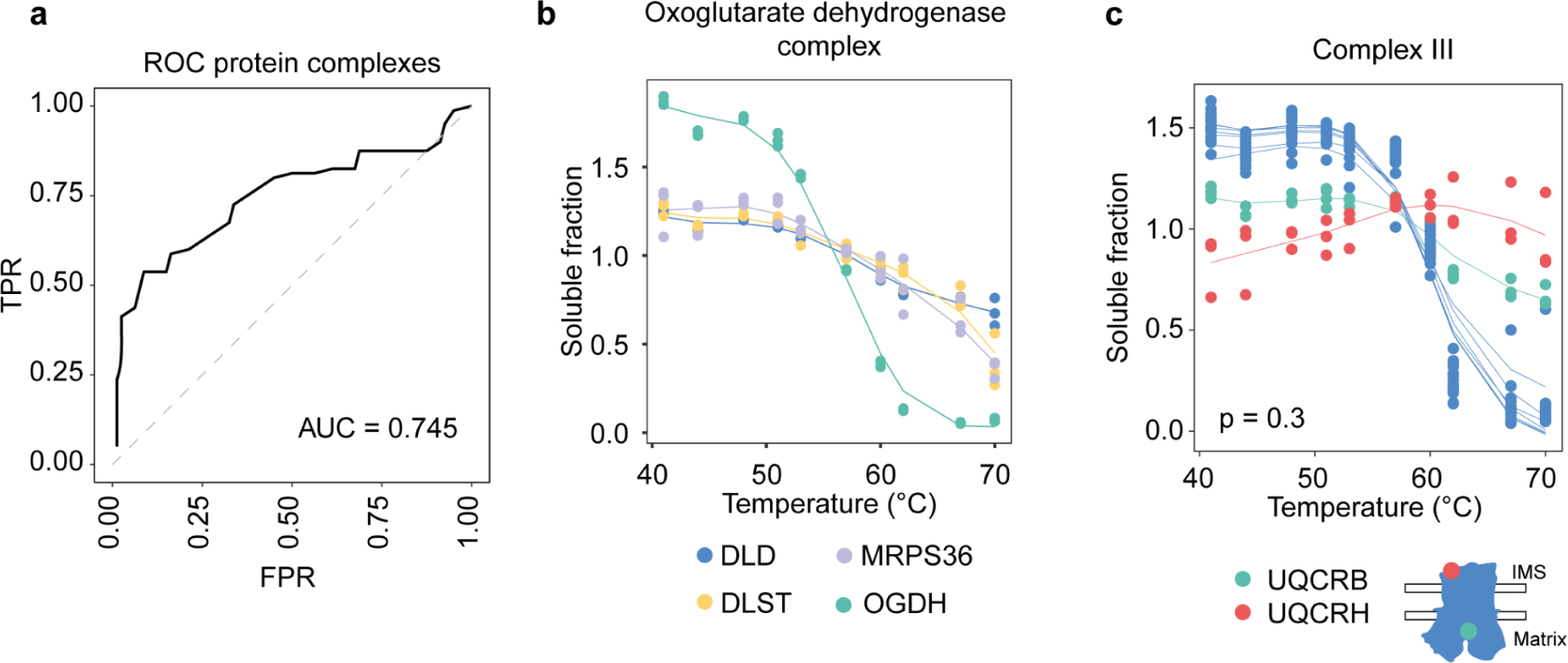
Performance of mitochondrial TPCA. **a**, Receiver operating characteristic (ROC) of recovered mitochondrial complexes (Supplementary Table 1). FPR = false positive rate; TPR = true positive rate; AUC = area under the curve. **b**, Melting curves of oxoglutarate dehydrogenase complex subunits. Data are fitted splines (df = 5). p = adjusted p-value (n = 3). **c**, Melting curves of complex III subunits. The draw represents the spatial distribution of the indicated proteins within the complex. Data are fitted splines (df = 5). p = adjusted p-value (n = 3).

**Supplementary Figure 6:**
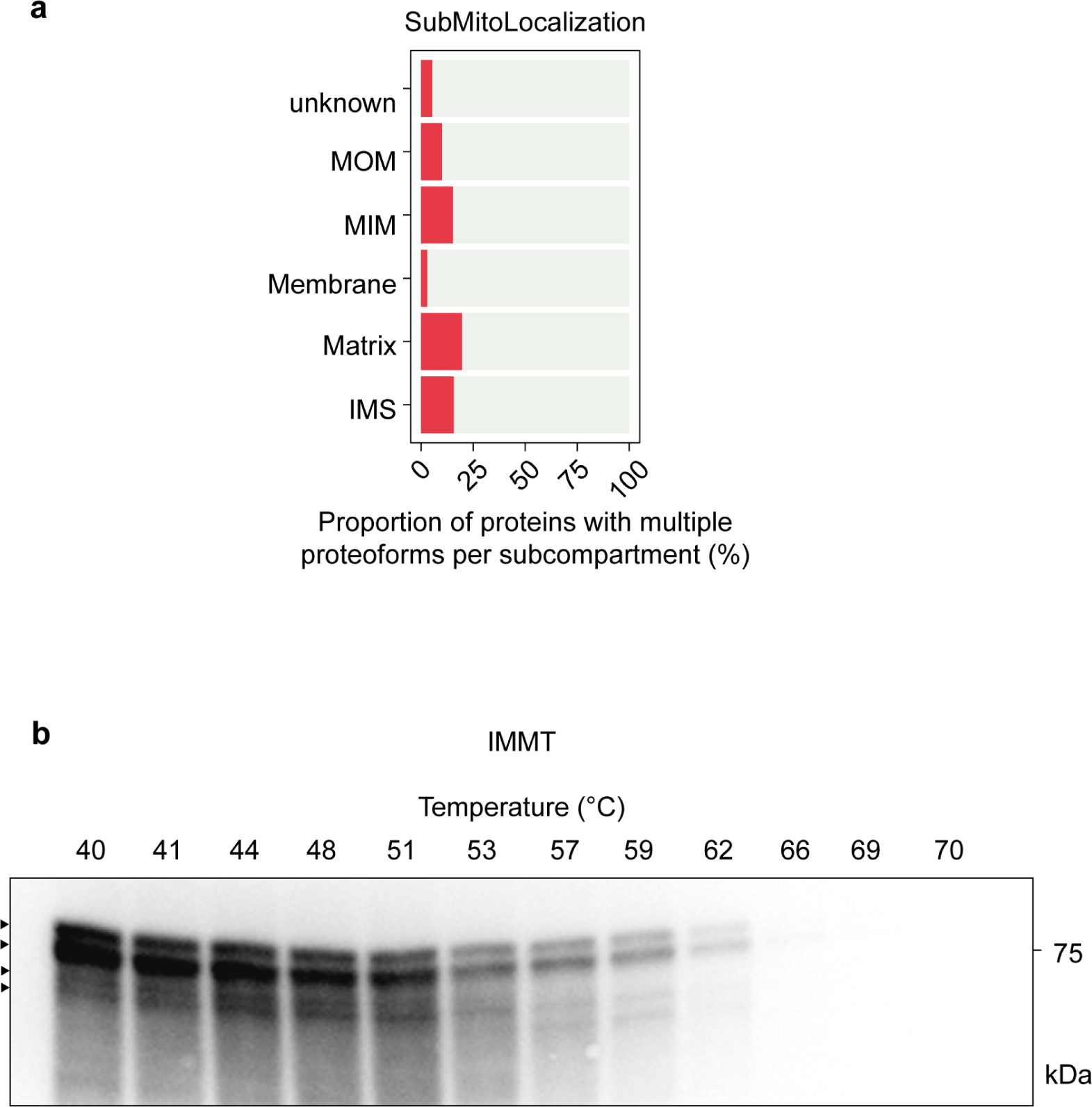
Exploration of mitochondrial proteoforms. **a**, Sub-mitochondrial localization of mitochondrial proteins with multiple proteoforms in proportion to all proteins annotated in that subcompartment. **b**, Immunoblot of the soluble fraction of IMMT from mitochondrial samples heated at the indicated temperatures (n=1). Arrows indicate the proposed proteoforms.

